# Older individuals do not show task specific variations in EEG band power and finger force coordination

**DOI:** 10.1101/2024.01.31.578126

**Authors:** Balasubramanian Eswari, Sivakumar Balasubramanian, Varadhan SKM

**Author notes:** Address correspondence to: Varadhan SKM, PhD, Department of Applied Mechanics and Biomedical Engineering, Chennai 600036, Tamilnadu, India.

## Abstract

**Background:** Controlling and coordinating finger force is crucial for performing everyday tasks and maintaining functional independence. Aging naturally weakens neural, muscular, and musculoskeletal systems, leading to compromised hand motor function. This decline reduces cortical activity, finger force control and coordination in older adults.

**Objective:** To examine the relationship between EEG band power and finger force coordination in older individuals and compare the results with young adults.

**Methods:** Twenty healthy young adults aged 20–30 (26.96±2.68) and fourteen older adults aged 58–72 (62.57±3.58) participated in this study. Participants held the instrumented handle gently for five seconds then lifted and held it for an additional five seconds in the two conditions: fixed (thumb platform secured) and free condition (thumb platform may slide on slider).

**Results:** In the older individuals there was no difference observed in the finger force synergy indices, and EEG beta band power between the two task conditions. However, in the young group synergy indices and EEG beta band power were less in free condition compared to fixed condition. Additionally, in the fixed condition, older adults showed a reduced synergy indices and reduced EEG beta band power than the young adults.

**Conclusion:** Older participants showed consistent synergy indices and beta band power across conditions, unlike young adults who adjusted strategies based on tasks. Young adults exhibited task-dependent finger force synergy indices contrasting with older individuals. These results suggest that the EEG beta band power may serve as a marker for finger force coordination rather than indicating the magnitude of force.

## Introduction

The ability to manipulate objects with the hand is often compromised in older individuals [1–3]. This decline in hand function is part of the multifaceted ageing process, which involves changes in both brain structure and functional anatomy [4,5]. The potential consequences of these changes include a broader deterioration in motor performance, coordination and balance as well as changes in gait [6]. During simple lift tasks, older adults tend to exert larger grip forces than younger adults [2]. The major cause of the decreased motor performances in older adults is due to the changes in brain structure including loss of grey matter [7], diminished dendritic density, reduced cerebral blood flow [8], diminished proprioception [9] and reduced peripheral nerve conduction [10]. Functional neural activity in motor-cortical areas can be evaluated by brain imaging systems like functional magnetic resonance imaging (fMRI), or the electrical activity of the cortical regions can be assessed by electroencephalogram (EEG) [11]. During gripping tasks, older adults exhibited diminished activation in the primary motor cortex (M1), contralateral primary sensory cortex (S1) and contralateral dorsal premotor cortex compared to the young adults [12]. In tasks involving thumb opposition, elbow flexion/extension and visually guided gripping, older adults exhibited higher activation in primary motor areas compared to young individuals. These indicate that the older individuals need more neural resources compared to the young individuals [12–14].

In EEG analysis, the alpha (8 – 12 Hz) and beta (13 – 30 Hz) frequency bands are considered to be associated with motor processes and the integration of sensory information [15]. During grip force control tasks such as force tracking tasks, older adults exhibit higher beta power and greater hemodynamic activity than young adults [16]. Additionally in simple motor tasks like finger pressing or tapping, older adults display a more pronounced decrease in alpha and beta band power over motor and parietal areas [5,17–19]. Furthermore, in older adults, the amplitude of movement-related beta band desynchronization was higher compared to that in the young group during dynamic and sustained handgrip paradigms [4]. Interestingly, during an elbow flexion task in older adults, EEG beta band power remains unchanged despite an increase in force [20].

Finger force synergies play a crucial role in object manipulation. In this context, we define synergy as a coordinated adjustment of finger forces and moments produced by each finger during multi-finger prehension tasks [21]. Synergy index is used to quantify the extent of co-variation among the forces and moment of forces produced by individual fingers, called elemental variables. It measures the degree to which the modulation of individual finger forces contribute to the reduction of variability in vital performance variables, like total grip/normal force, total moment of force, and total tangential/load force [22–24]. Several studies have provided evidence of age-related alterations in multi-digit synergies and the impaired coordination of finger forces during grip force modulation [25–28]. Grip force modulation relies on the continuous transformation of visual information into motor commands and the seamless incorporation of sensory response in the motor-cortical network [29].

Previous research has provided valuable insights into finger force control and the coordination of finger forces during object stabilization in young and older healthy populations. It consistently indicates that, in general, individuals over the age of 60 tend to apply greater grip force than young adults [2]. Despite this general trend, EEG analysis demonstrated that the specific neural processes and cortical activations associated with grip force among the older population can vary depending on the task being performed [16,20]. This implies that the association between the grip force coordination and neural activity varies across different activities. They are influenced by the nature of the task. Previous studies have explored EEG beta band power in the context of motor tasks, including finger tapping and isometric force production tasks [5,17,19].

Furthermore, earlier studies revealed that cortical activity does not directly control factors such as digit forces, net forces, positions of the fingers and the torque of the digits during object manipulation [30]. The synergy index serves as a quantitative measure that can be defined as the combined effect of both the central nervous system (CNS) and the intrinsic properties of the hand [31,32]. Synergy index quantifies the coordination of finger forces, not only reflect the mechanical properties but also the indicates the result of the neural modulation by the CNS. As individuals age, there might be less coordination and cooperation among finger forces [26] resulting in a lower synergy index. Hence, in this study the primary objective was to examine the relationship between finger force synergy index and EEG band power during lifting tasks. We hypothesized that older group would demonstrate lower finger force synergy indices. Additionally, we expected a greater EEG band power in the older group due to the age-related changes in the neural activity.

## Materials and Methods

### Participants

Twenty healthy right-handed young participants aged 20 to 30 years (26.95 ± 2.68) and fourteen healthy right-handed older participants aged 58 to 72 years (62.57 ± 3.58) were recruited for the study. None of the participants reported any neuromuscular disorder or injuries.

### Ethical approval

The experimental methods were approved by the Institutional Ethics Committee at the Indian Institute of Technology, Madras (Approval number: IEC/2021-01/SKM/02/05). Prior to start of the experiment, all participants provided written informed consent in accordance with the protocol adhering to the ethical guidelines of the IIT Madras. The experiments were conducted in adherence with the procedures approved by the Institutional Ethics Committee.

### Experimental Apparatus

A handle with five force sensors – Nano 17 (Force resolution: 0.0125 N, ATI Industrial Automation, NC, USA) was designed to measure the 6-component force/torque exerted by individual fingers, as shown in Figure 1 A. Four sensors were attached to one side of the designed handle to measure the four fingers I (index), M (middle), R (ring) and L (little) forces. One sensor to measure the force exerted by thumb was placed on a sliding platform opposite to the other four sensors. This arrangement allows vertical movement of the thumb with the option of fixing the thumb sensor anywhere on the handle. The inter-finger distance (center to center) between the index, middle, ring and little fingers is 20mm. The handle weighed approximately 0.750 kg, including an external weight of 0.250kg. The vertical movement of the thumb sensor was monitored using a displacement measurement sensor (OADM 12U6460, resolution 5 μm, Baumer, India). An acrylic block was positioned on the handle. The orientation of the handle was measured using an inertial measurement unit – IMU sensor (BNO055, with 16 bits resolution and operates with a range of 2000°/s, BOSCH, Germany), which was positioned on one side of the acrylic block. Orientation of the handle was measured to ensure that the handle did not tilt beyond a predefined angular limit so that there were no unaccounted torques in the system. A sprit level with bubble was also placed on the handle to enable the participants to visually focus and maintain the handle stable without tilting.

**Figure 1.**
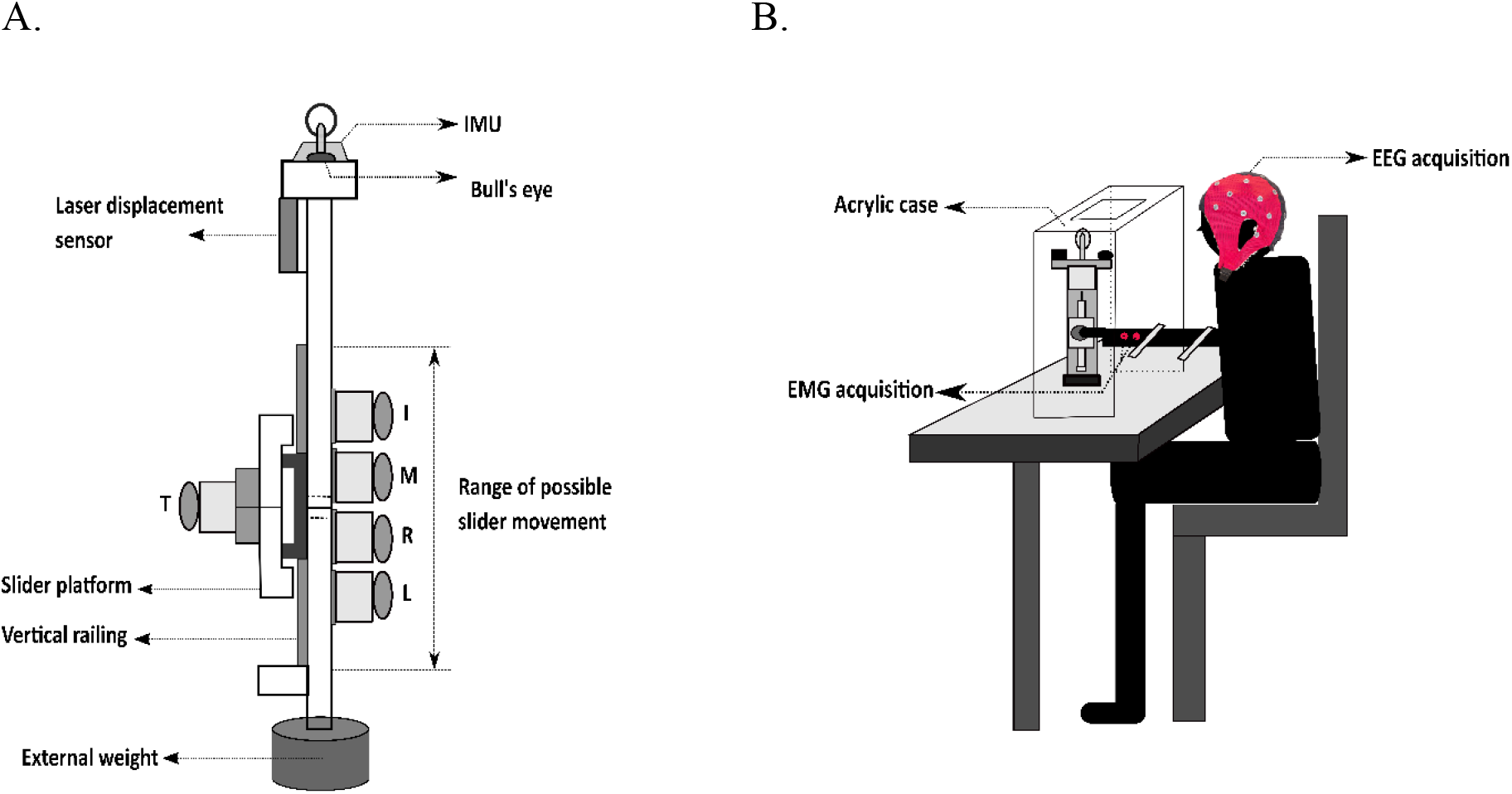
A) Instrumented Handle. The experimental handle, measuring 21 × 1 × 3 cm, consisted of an aluminium frame housing five force sensors, an IMU – for measuring the handle orientation and a displacement measurement sensor are shown. Five finger positions are denoted by the letters I (index), M (middle), R (ring), L (little), and T (Thumb). The handle had a net weight of 750 grams, included an external weight of 250 grams. **B) Experimental Setup**. The instrumented handle was placed inside an acrylic case. Participants were asked to gently grip and hold the handle for initial 5 seconds. Following this, participants were instructed to lift the handle and hold it in the air for 5 seconds.

Twenty-two channel EEG and two-channel EMG data was recorded with INTAN RHD 2216 amplifier ICs (Intan Technology, USA). RHD 2216 amplifiers are bipolar electrophysiology amplifier arrays with serial peripheral interface (SPI) and an on-chip analog-to-digital converter (16-bit ADC). 64-channel EEG cap (Waveguard original cap, 64-channel ANT neuro waveguard, The Netherlands; 10/10 electrode montage) was connected to two RHD amplifiers through a designed interface board. Each electrode from the EEG cap was connected to the RHD amplifiers. EEG signal was collected from the following 22 channels Fp1, Fp2, AFz, Fz, F3, F4, F7, F8, Fc1, Fc2, Fc5, Fc6, Cz, C3, C4, Cp5, Cp1, Cp2, Cp6, Pz, P3, P4, P7, P8, Oz, O1 and O2 according to the 10 – 10 electrode montage. The participants were instructed to wash and dry their hair prior to the experiment to achieve low impedance. EMG signal was collected form flexor digitorum superficialis muscle and flexor carpi ulnaris muscle. EMG analysis is not discussed and presented in this work.

### Experimental setup and protocol

For the experiment, the participants were required to sit on a chair and rest their right arm comfortably on the table, as shown in Figure 1 B. An EEG cap was placed on their head, and the electro-gel was applied to the electrodes using the syringe and blunt needle before the experiment. Electrode impedance was measured before the start of the experiment and set below 50K Ohms. The experiment involved of two task conditions: “fixed” and “free” thumb. The thumb slider which can move freely was fixed by a screw at the midpoint between the sensors of the middle and the ring finger under the fixed condition. Thumb slider had unrestricted movement as the screw was removed under the free condition. Participants were directed to keep the thumb platform positioned between the middle and ring fingers throughout the free task. The position of the thumb platform was measured using a displacement sensor. The trial was rejected if the thumb platform deviated more than 1cm above or below in the free condition.

In both fixed and free task conditions, initially, participants gently grasped the handle for 5 seconds in a relaxed manner. Following this, they were asked to lift the handle and sustain it in the air for 5 seconds using all fingers without any tilt. Throughout the 5 seconds of hold duration, participants were instructed to ensure the stable positioning of the handle by aligning the bubble level. Twenty-five such trials were taken for each participant for each condition. Sufficient breaks were given between trials, and breaks were provided upon the participant’s request.

### Data Acquisition

NI USB DAQ 6225 and 6210 with 16-bit resolution (National Instruments, Austin, TX, USA) were used to digitize thirty analog signals acquired from five force/torque sensors as well as the displacement signal from Baumer. These signals and orientation data from IMU sensors were sampled at 100 samples per second. Twenty-two channel EEG signals were collected using two RHD 2216 (Intan Technologies, USA) through XEM6310 (Opal Kelly, Portland) FPGA interface board. EEG signals were sampled at 1500 samples per second. Twenty-two channel EEG data, thirty force/torque signals, the displacement data, and the orientation were collected simultaneously using a custom LabView code (version 19).

### Data Analysis

All recorded signals were analyzed offline in MATLAB (Version R2022a, Math Works, USA).

### EEG Analysis

The power line interference and its harmonics from the recorded EEG signals were removed using a 50 Hz notch filter. Out of 22 channels, the following 14 EEG channels were taken for further analysis, Fc1, Fc2, Fc5, Fc6, Cz, C3, C4, Cp5, Cp1, Cp2, Cp6, Pz, P3, and P4. The notch filtered 14 channel EEG signals were bandpass filtered to the delta band (4 – 8 Hz), the alpha band (8 – 12 Hz), and the beta band (13 – 30 Hz) using a fourth order, zero phase lag Butterworth filter. Each band signal was down-sampled to 100 Hz, and the following steps were used to calculate the Band Power - BP [33] for both the fixed and free condition and the young and older group.

1. The power of the samples was obtained by squaring the amplitude samples
2. Averaging of the power samples across trials3.
3. Averaging of the samples over time to reduce the variability of the data

The band power value of the EEG signal between 2 – 3 seconds of the holding phase was taken to compare the two conditions for the 14 channels mentioned above. The mean band power during the holding phase was averaged across all participants, and the standard error of the mean was calculated.

### Synergy analysis

The covariation of digit force was calculated to observe the presence of synergy. Synergy is a set of elemental variables (EV) that covary to stabilize the performance variables (PV) [23]. Variance analysis was performed to quantify the multi-finger synergies [34–36]. An index of synergy (ΔV) was computed at the four digits (index, middle, ring, and little), known as virtual fingers–thumb level (VF – TH level), to measure the amount of covariation that occurs. The following are the equilibrium equations,

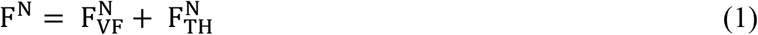

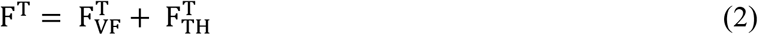

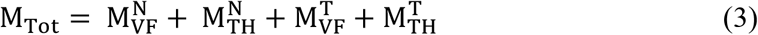

The superscripts T refers to the tangential force, and N refers to the normal force. The subscripts VF refers to the virtual finger, and TH refers to the thumb. M_Tot_ refers to the total moment. The performance variables variance was compared to the elemental variables variance and normalized by the elemental variables. The synergy index was computed using the following equations.

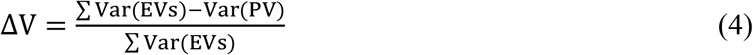

The synergy index was computed individually for each participant and condition. Positive values of the synergy index indicate the negative covariation among the elemental variables and indicate, a synergy stabilizing the performance variable. Negative values of the synergy index indicate an absence of synergy. Fisher’s z transformation was estimated to the synergy index using the equation (5) for the statistical analysis.

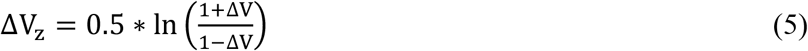

### Statistical analysis

The EEG band power and the behavioral data of the two conditions performed by 20 young and 14 older participants were considered for the statistical analysis. The analyses were executed using R. All the results are reported in the mean ± S.E.M format. A two–way repeated measures ANOVA was conducted with factors task conditions – 2 conditions: Fixed and Free X group – 2 groups: young and older to study the changes in synergy indices, EEG band power and thumb normal force. Tukey post-hoc test was also done for the comparison. The significant level p < 0.05 was chosen for the statistical analysis. Partial eta-squared (η^2^) was stated as effect size. The sphericity test was executed on the data for all the cases and Huynh-Feldt (H-F) criterion was applied as necessary to adjust the degree of freedom.

## Results

### EEG measures

EEG beta band power of the channel C3 of the young group in the fixed task condition was greater (Fixed condition: 0.064 ± 0.009 μV2, Free condition: 0.028 ± 0.004 μV2) than the free task condition (F (1,13) = 14.78, η2= 0.54, p < 0.05). Similarly, EEG beta band power of the young group was greater than the older group (Young group: 0.064 ± 0.008 μV2, Older group: 0.024 ± 0.005 μV2, in the fixed task condition (F (1,13) = 8.494, η2= 0.39, p < 0.05). No difference was observed in the EEG beta band power between the task condition in the older group as well as between the group in the free task condition, as shown in Figure 2.

**Figure 2.**
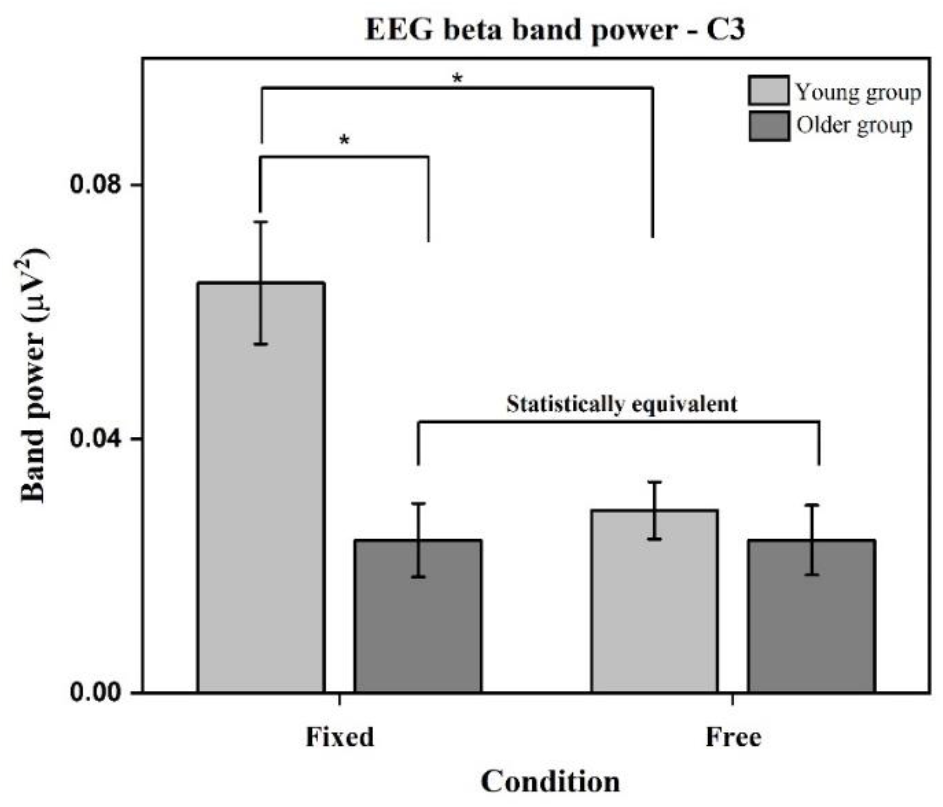
C3 channel EEG beta band power. In the older group, the beta band power of the channel C3 between two task conditions were statistically equivalent (t (13) = 0.012 – TOST analysis). However, in the young group, the beta band power of C3 channel was higher in fixed task condition compared to free task condition (p < 0.05). Additionally, beta band power of C3 channel of young group was higher compared to older group in the fixed task condition (p < 0.05).

Since there was no difference observed on the task condition in the older group, Two One-Sided Test (TOST) was performed to examine the equivalence between the conditions. TOST procedure was performed on the EEG beta band power comparing fixed and free task conditions within the older group. Equivalence bounds were estimated based on the SESOI = 1.04 (smallest effect size of interest). The lower and upper equivalence bounds were set as DL = -1.04 and DU = 1.04, respectively, for the statistical power of 95%. TOST procedure revealed a statistical equivalence on the EEG band power of the older group for the two task conditions (Fixed: 0.024 ± 0.006 μV^2^; Free: 0.024 ± 0.005 μV^2^, t (13) = 0.012, p < 0.05).

## Synergy Index

### Synergy index of the total moment of force

Synergy index of the total moment of force of the young group in the fixed task condition was greater (Fixed condition: 1.23 ± 0.13; Free condition: 0.38 ± 0.09) than the free task condition (F (1,13) = 14.43, η^2^ = 0.53, p < 0.05). Also, synergy index of the total moment of force of the young group was greater than the older group (Young group: 1.23 ± 0.13, Older group: 0.31 ± 0.14), in the fixed task condition (F (1,13) = 4.47, η^2^= 0.25, p < 0.05). No difference was observed in the synergy index of total moment of force between the task conditions in the older group as well as between the group in the free task condition, as shown in Figure 3.

**Figure 3.**
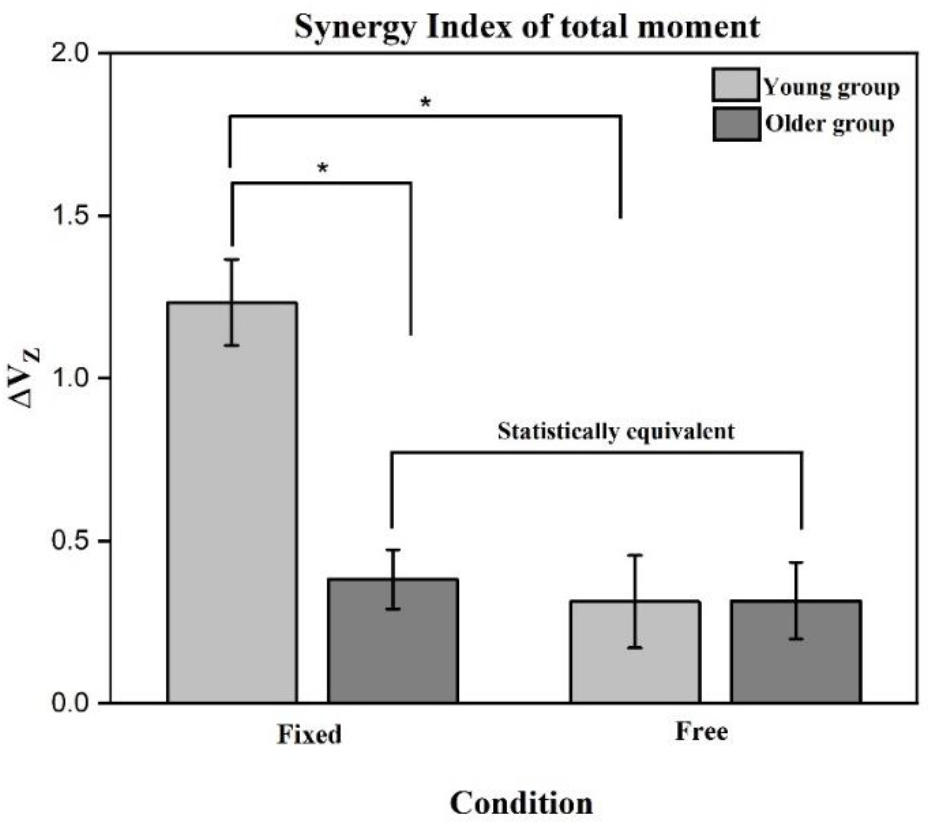
Synergy index of the total moment of force. In the older group, the synergy index of the total moment of force between the task conditions were statistically equivalent (t (13) = 9.11 – TOST analysis). However, in the young group, synergy index of the total moment of force in the fixed task condition was higher compared to the free task condition (p < 0.05). Furthermore, the synergy index of the total moment of force of the young group was higher than the older group in the fixed task condition (p < 0.05).

TOST (Two One-Sided Test) procedure was performed on the synergy index of the total moment of force between the fixed and free task conditions in the older group. From the SESOI calculations the upper and lower equivalence bounds were chosen as DU = 1.04 and DL = -1.04, respectively, for the statistical power of 95%. TOST procedure revealed a statistical equivalence (t (13) = 9.11, p < 0.05) on the synergy index of the total moment of force of the older group for the two task conditions (Fixed condition: 0.31 ± 0.14; Free condition: 0.32 ± 0.12).

### Synergy index of total tangential force

Synergy index of the total tangential force of the young group in the fixed task condition was higher (Fixed condition: 1.97 ± 0.28; Free condition: 0.56 ± 0.14) than the free task condition (F (1,13) = 31.57, η^2^ = 0.71, p < 0.05). Also, synergy index of the total tangential force in the fixed task condition of the young group was greater (Young group: 1.97 ± 0.28; Older group: 0.58 ± 0.12) than the older group (F (1,13) = 6.45, η^2^ = 0.33, p < 0.05). There was no difference observed between the fixed and free task conditions in the older group as well as between the group in the free task condition as shown in Figure 4.

**Figure 4.**
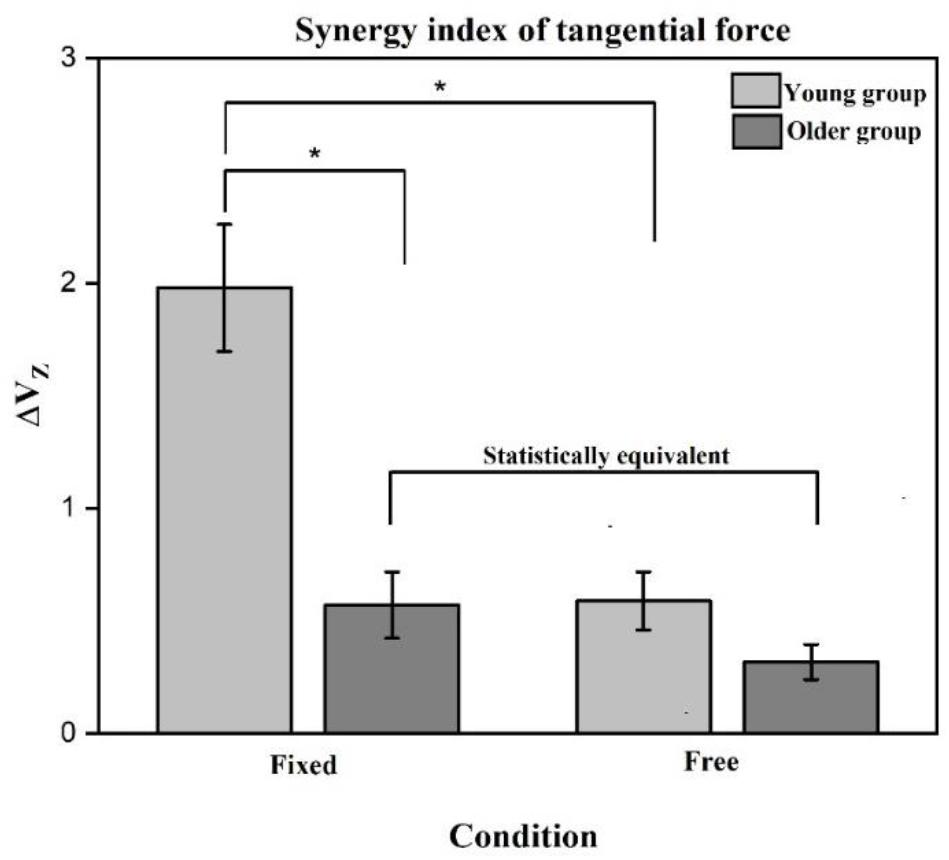
Synergy index of total tangential force. In the older group, the synergy index of the total tangential force between the task conditions were statistically equivalent (t (13) = 9.33 – TOST analysis). However, in the young group, synergy index of the total tangential force in the fixed task condition was higher compared to the free task condition (p < 0.05). Furthermore, the synergy index of the total tangential force of the young group was higher than the older group in the fixed task condition (p < 0.05).

TOST (Two One-Sided Test) procedure was performed on the synergy index of total tangential force between the fixed and free task conditions in the older group. With the small effect size of interest (SESOI) calculations the upper and lower equivalence bounds were chosen as DU = 1.04 and DL = -1.04, respectively, for the statistical power of 95%. TOST procedure revealed a statistical equivalence (t (13) = 9.33, p < 0.05) on the synergy index of the total tangential force of the older group between the two task conditions (Fixed condition: 0.58 ± 0.12; Free condition: 0.32 ± 0.29).

### Discussion

The primary aim of the current study was to investigate the relationship between the EEG band power and finger force synergies during object manipulation in the healthy young and older participants. It was hypothesised that the older individuals would demonstrate lower finger force synergy and higher EEG band power. As expected, the older group demonstrated lower synergy indices indicating diminished finger force coordination. Contrary to the hypothesis beta band power measured from the C3 channel of the EEG was found to be lower in the older participants than the young participants. Additionally, in the older participants, synergy index of tangential force, synergy index of the total moment of force, and the EEG beta power showed no noticeable difference between the two task conditions. In contrast, the synergy indices and beta band power of the young were found to be considerably lower in the free condition than the fixed condition. These findings imply that the young participants use different strategies in the two tasks whereas older individuals exhibit a consistent pattern of behaviour across tasks. Furthermore, these findings indicate that cortical activity may correlate with synergistic activity rather than solely influencing the finger force.

The reduction in beta band power in the older individuals could be attributed to the age-related functional deficits resulting in reduced muscle strength. The beta band activity reflects the synchronous firing of neurons group in the motor cortex during sustained muscle contractions [37]. The reduction in beta band power within the older group might be due to the age-related changes in neural synchronization. The changes in the neural synchronisation may lead to the alterations in the efficiency of information processing between brain regions. Though there is a decrease in band power, the older individuals could lift the handle in both task conditions. This may be due to the recruitment of neurons from other subcortical regions to finish the task, suggesting compensatory mechanisms invoked by the aging brain [5]. The young participants exhibited considerably higher EEG band power than older participants in the fixed task condition. This is in line with the previous findings which showed a decrease beta band activity in the older individuals during force tracking tasks [16].

EEG band power and the synergy indices of the young group is higher in the fixed condition compared to the free task condition. In the free condition, friction between the platform and the thumb is changed which can be perceived by the proprioceptors and the cutaneous receptors of the fingertip. Handle’s equilibrium was balanced by adjusting the tangential and normal forces of the fingers. Notably, among young individuals, a substantial reduction in the synergy indices and EEG band power was observed in the free condition in contrast to the fixed condition. These results suggest that the capability to stabilize critical performance variables through the covariation of elemental variables is reduced in the free condition, especially in the young individuals. This implies a greater need of task specific finger force adjustments in synergy in the free task condition possibly attributed to the friction conditions. Reduction in synergy indices can be seen as reduced coordination of finger force in the free condition. Such adjustments in the finger force synergy may be attributed to the response of the tactile feedback from the neural system and the implementation of specific strategy in the young individuals. Additionally, EEG band power of the free condition is also considerably less in young individuals compared to the fixed task condition. It is known that when muscles work together in a synergistic manner, their coordinated activity is accompanied by a shared cortical pattern in beta band frequency [38,39]. The shared cortical pattern suggests a synchronised firing of neurons in the brains motor areas indicating high degree of coordination among the finger muscles. Indeed, this coordinated muscle activity can be seen as functional unit or synergy at the cortical level [40,41].

In the fixed task condition, there is a higher synergy in the young group. The increased synergy indicates the more effective and coordination which is associated with higher EEG band power. This can be seen as that the enhanced force synergy is associated with stronger synchronisation and activation of neural networks involved. On the contrary decreased synergy or less coordination has been associated with less band power which could be seen in the free task condition in the young group. It is also noted from the previous studies that the EEG – EMG coherence is reduced when there is a need for individuated muscle control. These findings support the idea that cortical network activity plays a critical role in coordinating synergistic control rather than solely determining the magnitude of the finger force. This is in agreement with the findings that EEG features may not predict the magnitude of the force. [42]. The synergy indices of the older individuals were substantially less than the young group in the fixed task condition, which is in agreement with previous findings [23]. This indicates that older individuals have reduced coordination and integration of finger forces.

Aging is associated with the diminished tactile sensations [2,43,44] and a reduced ability to process sensory feedback [3]. The reduction in sensory feedback affects the CNS control in the older group leading to a decline in the adaptation of finger force coordination or high individuation of finger force as reflected in reduced synergy indices. This decrease in adaptation manifests as a uniform or consistent pattern to meet the task demands. These findings also indicate diminished finger force coordination in the older group compared to young individuals. The results indicate that older participants exhibit a uniform finger force coordination pattern across different tasks. However, these findings appear to contradict earlier reports indicating high synergy indices during rotational task movements in the older individuals [28].

In the older individuals EEG band power was less compared to the young participants. These results contradict earlier research where the EEG beta band power of the older participants was higher compared to the young participants in the elbow flexion task [18]. This could be due to the requirement of the higher force in the elbow flexion tasks. The findings suggests that finger force coordination and integration may differ depending on task demands among the older group. However, such a difference could also be attributed to the differences in the cortical control of distal and proximal segments. The distal segments have been observed to have greater monosynaptic projections from the cortical neurons compared to the proximal segments. Also, no noticeable difference was observed in the EEG beta band power of older adults between the two-task conditions. This lack of difference is also observed in the synergy indices, suggesting consistent performance levels across different conditions in the older participants. The lack of difference was statistically equivalent as supported by the TOST statistical analysis.

Taken together, the older individuals exhibited diminished capacity in finger force coordination and cortical network activity compared to the young participants, is in accordance with the earlier findings [45]. This can be interpreted as young individuals employed diverse strategies to regulate the coordination of finger force as evidenced by variations in the synergy indices and the EEG band power across tasks. On the contrary, older individuals seem to rely on a more uniform or consistent strategy across tasks evident both in the finger force synergy indices as well as in the EEG band power. This could be attributed to a diminished or loss of sensory information in the older individuals particularly in addressing the changes in the task conditions. The decrease in band power and synergy indices in older individuals could arise not only from factors such as reduced muscle strength and diminished motor unit phenomena at the peripheral level. But also, changes in the cortical level and strategic control adjustments contribute to these uniform patterns in the older individuals. Previous studies documented that the reduced tactile sensation is the main cause of the increased grip force in older individuals [1–3]. Consequently, this sensory decline may be a key factor preventing the older individuals from effectively modulating finger force synergy indices in response to varying task conditions.

These findings align with previous research in certain aspects [20,26] but differ in others [28], suggesting that the coordination patterns may vary depending on the task requirements. There is a lack of consistent conclusions regarding the effects of hand postures and finger force on EEG activity. Various studies have reported different findings, contributing to the complexity of understanding the relationship between EEG band power and motor tasks. Different force loads did not affect EEG band power, suggesting that force levels may not impact EEG activity [46]. These discrepancies in EEG findings highlight the complex relationship between EEG beta band power and hand activities, which may depend on factors such as experimental protocols, cortical regions of interest, and algorithms for data analysis. These factors can significantly influence the observed results and make it challenging to draw consistent conclusions. Further research is required to explore the above-mentioned discrepancies and understand the underlying mechanisms. By considering multiple factors and adopting different protocols, future studies can provide more comprehensive insights into the relationship between EEG activity and kinetics during object manipulation.

### Conclusion

This study examined the neural signatures of finger force synergy in healthy young and older individuals. The results indicated that the EEG beta band power was substantially higher in young participants than in older participants in fixed task condition, consistent with previous research. Additionally, the young participants demonstrated a reduction in EEG beta band power and synergy indices, demonstrating the task specific finger force control depending on the task requirement. Alternatively, the older participants did not exhibit changes in synergy indices, or EEG beta band power across different task conditions, indicating a limited ability to adjust synergies based on task demand. These findings suggest that the aging process influences the neural mechanisms of finger force coordination. The inability to adjust finger force synergy indices and beta band power among the older individuals serves a distinct indicator of age-related transformations in motor control.

## Supporting information

Supplementary results

