## Supplementary results for "Older individuals do not show task specific variations in EEG band power and finger force coordination"

**Supplementary material**

**Force Analysis**

Force signals recorded from the five sensors and the displacement data were applied through a second-order, zero-phase lag lowpass Butterworth filter with the cut of frequency of 15 Hz. The samples in the hold phase between 2 to 3 seconds were analyzed for each trial. The normal and tangential force for each fingertip force were averaged over the time samples, trials, and across participants, and the standard error of the mean was calculated.

**Normal forces and total tangential forces**

Thumb normal force of the young in the free task condition (Fixed condition: 9.69 ± 0.48; Free condition: 12.12 ± 0.44) was greater than the fixed task condition ($F_{(1,13)} = 15.47$, $\eta^2 = 0.54$, $p < 0.05$). Similarly, the thumb normal force of the older group was greater (Young group: 9.69 ± 0.48; Older group: 13.49 ± 0.65) than the young group in the fixed task condition ($F_{(1,13)} = 9.96$, $\eta^2 = 0.43$, $p < 0.05$). Additionally, no difference was observed between the groups in the free-task condition, and between the task condition in the older group as shown in Figure 1. The thumb normal force of older people in the fixed and free conditions showed no significant difference. TOST (Two One-Sided Test) procedure was performed on the thumb normal force dependent samples between the fixed and free task condition in the older group. The upper and lower equivalence bounds were chosen as $\Delta U = 1.04$ and $\Delta L = -1.04$, respectively, for the statistical power of 95% [47]. TOST procedure revealed a statistical equivalence ($t(13) = 3.15$, $p < 0.05$) on the thumb grip force of the older group for the two task conditions (Fixed: 13.49 ± 0.65; Free: 13.25 ± 0.65).

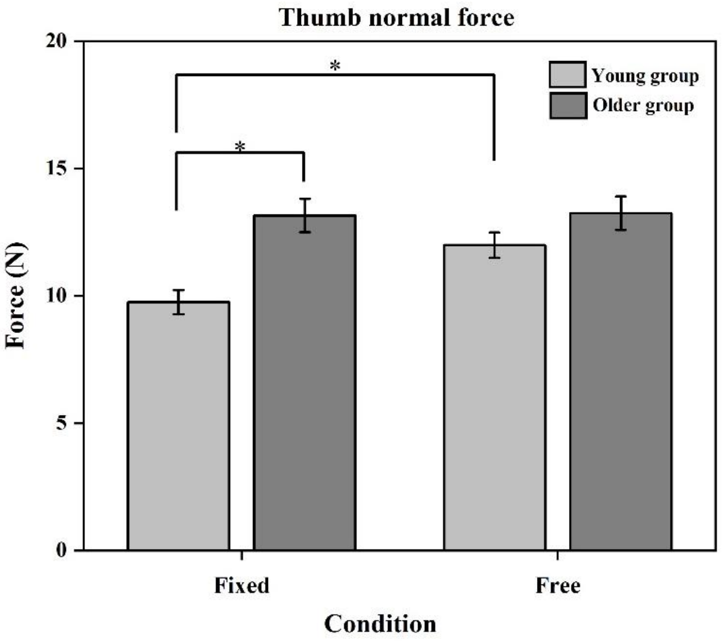


**Figure 1 Thumb normal force**. In older group, thumb normal force between the two-task condition were statistically equivalent (t (13) = 3.15, p < 0.05 – TOST analysis). However, in the young group, thumb normal force of the fixed condition was higher than the free condition (p < 0.05). Also, the thumb normal force of the older group was greater than the young group in the fixed task condition (p < 0.05).

There was no statistical difference observed between the conditions and group in the total tangential force. Total tangential force was maintained enough to lift and hold the handle (Young: Fixed condition: 7.67 ± 0.17; Free condition: 7.65N ± 0.13; Elderly: Fixed condition: 7.46 ± 0.28; Free condition: 7.32N ± 0.15).
